# Naturalistic audio-movies reveal common spatial organization across “visual” cortices of different blind individuals

**DOI:** 10.1101/2021.04.01.438106

**Authors:** Elizabeth Musz, Rita Loiotile, Janice Chen, Marina Bedny

## Abstract

Occipital cortices of different sighted people contain analogous maps of visual information (e.g., foveal vs. peripheral space). In congenital blindness, “visual” cortices enhance responses to nonvisual stimuli. Do deafferented visual cortices of different blind people represent common informational maps? We leverage a naturalistic stimulus paradigm and inter-subject pattern similarity analysis to address this question. Blindfolded sighted (S, n=22) and congenitally blind (CB, n=22) participants listened to three auditory excerpts from movies; a naturalistic spoken narrative; and matched degraded auditory stimuli (i.e., shuffled sentences and backwards speech) while undergoing fMRI scanning. In a parcel-based whole brain analysis, we measured the spatial activity patterns evoked by each unique, ten-second segment of each auditory clip. We then compared each subject’s spatial pattern to that of all other subjects in the same group (CB or S) within and across segments. In both blind and sighted groups, segments of meaningful auditory stimuli produced distinctive patterns of activity that were shared across individuals. Crucially, only in the CB group, this segment-specific, cross-subject pattern similarity effect emerged in visual cortex, but only for meaningful naturalistic stimuli and not backwards speech. These results suggest that spatial activity patterns within deafferented visual cortices encode meaningful, segment-level information contained in naturalistic auditory stimuli, and that these representations are spatially organized in a similar fashion across blind individuals.

**Significance Statement:** Recent neuroimaging studies show that the so-called “visual” cortices activate during non-visual tasks in people who are born blind. Do the visual cortices of people who are born blind develop similar representational maps? While congenitally blind individuals listened to naturalistic auditory stimuli (i.e., sound clips from movies), distinct timepoints within each stimulus elicited unique spatial activity patterns in visual cortex, and these patterns were shared across different people. These findings suggest that in blindness, the visual cortices encode meaningful information embedded in naturalistic auditory signals in a spatially distributed manner, and that a common representational map can emerge in visual cortex independent of visual experience.

## Introduction

Cognitive and perceptual processes localize to similar anatomical areas across people, as observed in retinotopic maps in V1 (e.g., Engel et al., 1997; DeYoe et al. 1996, Sereno et al. 1995). How do innate constraints and experience interact to produce these common informational maps? One way to gain insight into this question is to compare cortical organization across populations with different developmental experiences: people with and without vision.

Previous research finds that blindness enhances “visual” cortex responses during auditory, tactile and higher-cognitive tasks, such as Braille reading, sound localization, and spoken language comprehension (Sadato et al., 1996; Collignon et al., 2011, Roder et al., 2002; Bedny et al., 2011; Kanjlia et al., 2016). Do visual cortices of people born blind develop similar representational maps across individuals? Alternatively, do such common maps only emerge when visual cortices assume their evolutionarily predisposed visual functions?

Most neuroimaging studies of visual cortex plasticity in blindness have targeted a single cognitive process (e.g., sound localization, language) using simplified experimental designs (e.g., Poirer et al., 2006; Thaler et al., 2011). A handful of studies have identified anatomically separable responses to two or three information types within “visual” cortices of blind people (e.g., language vs. math) (Kanjlia et al., 2016; Abboud et al, 2019). Recent studies find that the ventral occipitotemporal and early visual cortices of people born blind show spatially distinct responses to stimuli from different semantic categories (e.g., faces and places) when these are presented either via touch or auditorily (van den Hurk et al., 2017; Mattioni et al., 2020; Ratan et al., 2020; Vetter et al., 2020; Pietrini et al., 2004). How consistent are such information “maps” across blind individuals?

We test this question by directly comparing cortical maps across blind and sighted people using shared pattern analysis. Recently developed fMRI analysis techniques provide new ways for directly studying shared information maps across brains by comparing multi-voxel activity patterns elicited by naturalistic stimuli across individuals. Naturalistic sounds, such as movie excerpts and spoken narratives, contain a range of rich information, from low-level perceptual to higher order semantic features. Each segment of an unfolding narrative induces a characteristic spatial pattern of activity that is similar across people and distinct from other segments in the narrative (Chen et al., 2017; Zadbood et al., 2017; Baldassano et al., 2018). In sighted adults, naturalistic stimuli induce shared patterns in higher-order cognitive networks implicated in semantic processing, including the precuneus and bilateral frontal and lateral temporal regions. Two recent studies showed that naturalistic auditory stimuli drive synchronized activity across “visual” cortices of blind individuals (Van Ackeren et al., 2018; Loiotile et al., 2019). However, whether sub-segments of these stimuli produce consistent spatial patterns across blind individuals has not been tested.

In the current study, congenitally blind and sighted blindfolded adults listened to auditory excerpts from popular live-action movies and a spoken narrative while undergoing fMRI. Participants also heard two control stimuli: one in which the narrative sentences were shuffled to disrupt the plotline, and the narrative played in reverse (backwards speech), removing semantic and linguistic information. We measured spatial activity patterns evoked by each ten-second segment of the meaningful (movies and narrative) and control (shuffled sentences and backwards speech) stimuli and used inter-subject spatial correlation analyses to test for shared, segment-specific patterns among individuals within each group. We predicted that the meaningful stimuli, but not controls, would reveal consistent information maps across blind people in visual cortex.

## Methods

### Participants

Twenty-two congenitally blind (CB, 8 males) and twenty-two sighted (S, 4 males) individuals contributed data to this study. All blind participants reported having minimal or no light perception since birth, such that they could not perceive motion, colors, or shapes. Sighted participants reported normal or corrected-to-normal vision. Groups were matched on average age (CB mean = 43.5, s.d.= 16.9; S mean= 42.7, s.d. = 13.8, *t*(21)= -0.17, *p*>0.5) and years of education (CB mean= 16.9, s.d.= 2.6; S mean= 18.1, s.d.= 2.62, *t*(21)= 1.2, *p*= 0.2). Data from one additional blind participant was excluded because after the scan they reported temporarily having vision during childhood. At the time of the study, participants were not taking psychoactive medications, by self-report, and reported no history of neurological disorders, brain damage, or head injuries. For blind participants, all causes of blindness were due to damage in areas posterior to the optic chiasm, such as the eye or optic nerve, and not brain damage (Table 1). All participants provided informed consent as approved by the Johns Hopkins Institutional Review Board. These data have been reported in previous analyses (cf. Loiotile et al., 2019).

**Table 1.** Participant count by etiology of blindness

| **Etiology** | **Participant number** |
| --- | --- |
| Leber congenital amaurosis | 7 |
| Retinopathy of Prematurity | 8 |
| Optic nerve hypoplasia | 3 |
| Retinitis pigmentosa | 1 |
| Unknown | 3 |

**Table 2.** Description of the main auditory stimulus conditions

| Stimulus Name | Condition | Duration (min) | TRs | RMS amplitude | Frequency (Hz) | Blind Participants | Sighted Participants |
| --- | --- | --- | --- | --- | --- | --- | --- |
| Rest | Non-intact | 7.4 | 240 |  |  | 22 | 21 |
| Backwards Speech | Non-intact | 6.8 | 223 | 0.032 | 1177 | 22 | 22 |
| Sentence-Scramble | Non-intact | 6.8 | 223 | 0.032 | 1177 | 22 | 22 |
| Pie-Man | Intact | 6.8 | 223 | 0.032 | 1177 | 22 | 22 |
| Movie 1 (*The Conjuring*) | Intact | 5.1 | 172 | 0.11 | 2472 | 22 | 22 |
| Movie 2 (*Taken*) | Intact | 5 | 169 | 0.031 | 1600 | 22 | 21 |
| Movie 3 (*Blow-Out*) | Intact | 6.5 | 215 | 0.054 | 4414 | 22 | 22 |

### Stimuli

The main testing materials consisted of six audio clips each lasting five to six minutes. Three of the audio clips were excerpts from popular live-action movies. These sound clips were selected to facilitate a shared interpretation and experience across participants, because they are suspenseful, engaging, and easy to follow. The remaining three audio clips were each created from the same six-minute excerpt of a live comedy sketch, “Pie-Man.” The intact version of Pie-Man was presented in its original format. The “Sentence-Shuffle” version was created by splicing and re-ordering the sentences of the intact version, such that the clip contained intelligible sentences but lacked a coherent plotline. Finally, the “Backwards Speech” version was created by playing the clip in time-reverse format, such that it lacked any intelligible content (cf. Loiotile et al., 2019). The intact, scrambled, and time-reversed versions of “Pie-Man” have been used in several previous neuroimaging studies to probe shared brain responses across sighted individuals (e.g., Lerner et al., 2011, Stephens et al., 2013; Simony et al., 2016). We also collected a resting state scan, during which no auditory stimuli were presented and participants were instructed to relax and stay awake. All stimuli are available for download at the Open Science Framework: https://osf.io/r4tgb/

### Procedure

Participants passively listened to each sound clip during fMRI scanning. Each clip was assigned to a separate scanning run and clips were presented in a pseudo-random order to each participant, such that the sequence for each blind participant was yoked to a corresponding sighted participant. For each “intact” stimulus, participants were read a 2-3 sentence prologue that provided context for the upcoming sound clip immediately before the scanning run began. Participants were instructed to listen carefully and pay attention, as they would later be asked questions about the story content of each clip. The presentation of each auditory clip was preceded by five seconds of rest and followed by 20-22 seconds of rest. These rest periods were not included in the analyses. All participants wore light exclusion blindfolds during scanning. All participants completed all seven scanning runs, except for one sighted participant who did not complete one audio-movie scanning run and the resting state scan due to time constraints.

Prior to scanning, we confirmed that each participant was not already familiar with the intact clips (i.e., they had not previously seen the movies that the clips were sourced from). After completing MRI scans, participants exited the scanner and immediately answered five orally presented multiple choice questions for each of the intact auditory clips. Although we included data from all participants in all reported brain analyses, results are qualitatively similar when analyses are limited to data from participants who scored 3/5 or better on all four post-scan comprehension tests.

Auditory stimuli were presented over Sensimetrics MRI-compatible earphones (http://www.sens.com/products/model-s14/) at the maximum comfortable volume for each participant. Prior to the main stimulus presentation and during the acquisition of the structural scans, we tested participants’ ability to hear sounds over the scanner noise. Volume levels were adjusted to maximize audibility and comfort (cf. Loiotile et al., 2019).

### fMRI Data Acquisition

Structural and functional MRI data of the whole brain were collected on a 3 Tesla Phillips scanner. T1-weighted structural images were collected in 150 axial slices with 1 mm isotropic voxels using a magnetisation-prepared rapid gradient-echo (MPRAGE). T2*-weighted functional images were collected with a gradient echo planar imaging sequences in 36 sequential ascending axial slices with 2.4 × 2.4 × 3 mm voxels and 2-second TR (echo time: 3ms, flip angle: 70 degrees, field-of-view: 76×70 matrix, slice thickness: 2.5mm, inter-slice gap: 0.5, slice-coverage: FH 107.5, PH direction L/R, first-order shimming).

### fMRI Preprocessing

Preprocessing was performed using FEAT (fMRI Expert Analysis Tool) Version 6.00, part of FSL (http://fsl.fmrib.ox.ac.uk/fsl) and included slice time correction using Fourier-space time-series phase-shifting, motion correction using MCFLIRT (Jenkinson 2002), high-pass filtering (140s), and linear detrending. All functional volumes were co-registered and affine transformed to a standard anatomical brain (MNI152) using FLIRT. Functional data were smoothed with a 4mm FWHM Gaussian kernel and resampled to 3mm isotropic voxels. The first four and last eight TRs of each scanning run were discarded, corresponding to the rest periods before and after the clips (accounting for delays in hemodynamic response). Analyses were performed in volume space and data were projected onto a cortical surface with the HCP Workbench for visualization (Marcus et al., 2011).

### Inter-Subject Pattern Similarity Analysis

We computed spatial pattern inter-subject correlations (spatial ISC) for each subject group (blind and sighted) and condition (Audio-Movies, Pie-Man, Sentence-Shuffle and Backwards Speech) (Chen et al., 2017; Nastase et al., 2019; Zuo et al., 2020). Pattern analysis was conducted in each of 400 parcels from an independent whole-brain resting-state parcellation (Figure 1C) (Schaefer et al., 2018). For each run, timeseries data were divided into a set of non-overlapping ten-second segments (32-47 segments per run) and averaged across TRs within a segment, yielding one voxel pattern of brain activity for each segment in each parcel (Fig. 1A; see Fig S1 for example segment-level patterns). Within each parcel, the Pearson correlation was calculated between every segment pattern for each participant versus every segment pattern averaged across the remaining participants, yield a segment-by-segment correlation matrix (Fig. 1B). The spatial ISC value is computed by averaging together the on-diagonal (i.e., matching) segment pattern similarity values. For the Audio-Movies condition, data from all three audio-movie clips were concatenated together prior to computing spatial ISC. The Audio-Movies analysis was limited to participants who had useable data for all three movie clips (CB n= 22, S n= 21). Results for each individual audio-movie stimulus are presented in supplementary materials.

**Figure 1.**
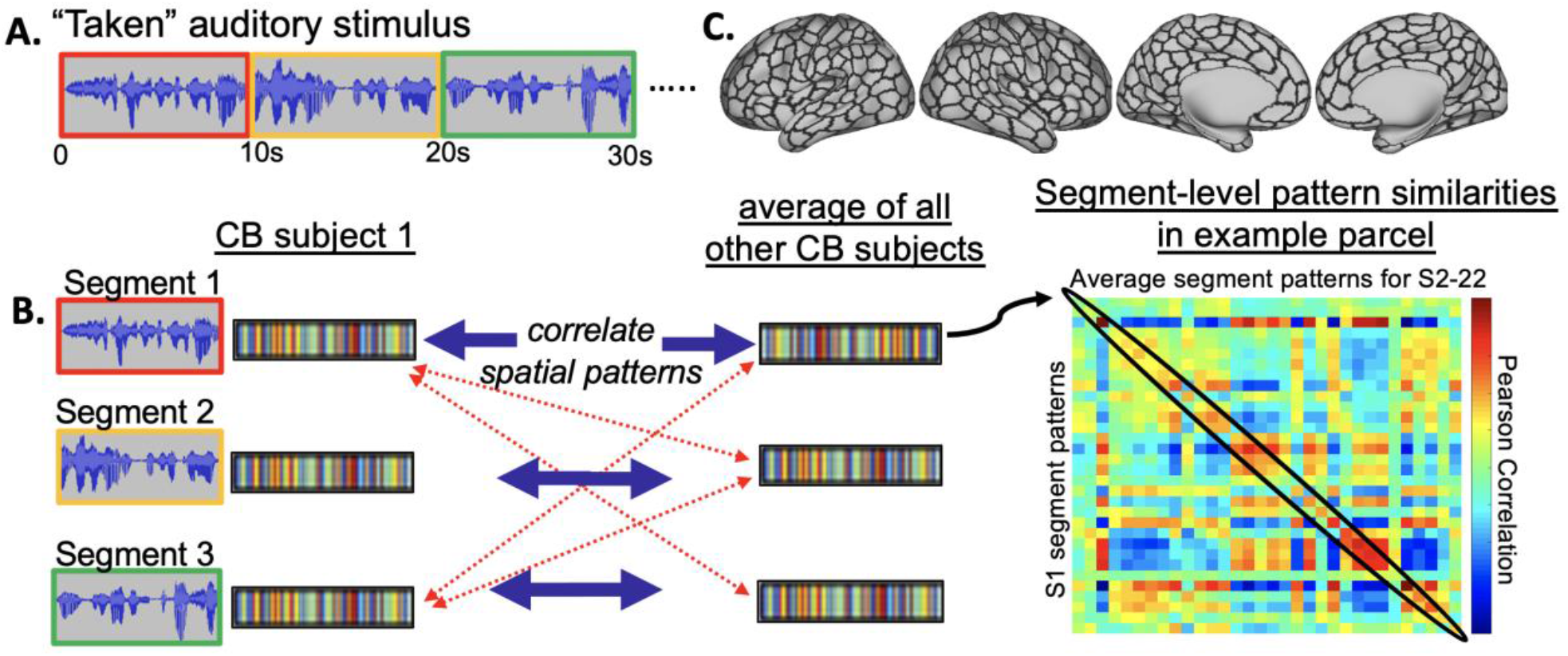
Analysis schematic of spatial pattern inter-subject correlation (ISC) analysis. A. Each stimulus was segmented into ten-second-long segments. B. Similarities were computed between one subject’s segment-level spatial patterns (S1) and the group-average patterns from the remaining members of their respective group (S2-22). The mean on-diagonal segment pattern similarity (circled) provides the spatial ISC value. C. Group-average spatial ISC values were computed in each of 400 parcels for each auditory stimulus. Parcellation was determined by a resting-state analysis on independent data (Schaeffer et al., 2018). CB = congenitally blind

Group-level spatial ISC values were computed by averaging the ISC values across segments and across participants. To determine statistical significance, the observed group-level spatial ISC values were compared to permutation-based null distributions. For each parcel, a null distribution was generated by randomly shuffling the spatial ISC values in the group-average segment-by-segment pattern correlation matrix (Fig. 1B) 1,000 times and retaining the group-average mean on-diagonal value of each resulting similarity matrix (Kriegeskorte, 2008). This resulted in 1,000 parcel ISC maps derived from scrambled spatial ISC values. From each null parcel map generated from this permuted data, the largest of the 400 spatial ISC parcel values was retained, yielding a null distribution of 1,000 maximum noise correlation values (cf. Regev et al., 2013). Family-wise error rate (FWER) was then defined as the top 5% of the null distribution of the maximum correlation values, which was used to threshold the observed parcel map (Nichols & Holmes, 2002). This analysis ensures that the patterns in above-threshold parcels contain segment-specific information, because the average correlation between spatial patterns for matching segments must exceed an equal-sized random sample of correlations between all segments (i.e., both matching and nonmatching) in order to meet statistical significance (cf. Chen et al., 2017).

#### Between-Condition Comparisons

To identify parcels where spatial ISC was reliably greater for one stimulus condition versus another, we computed a condition difference score for each participant (e.g., Audio-Movies spatial ISC minus Backwards Speech spatial ISC) and then averaged these scores across participants. To generate a null distribution, at each parcel we randomly flipped the sign on the difference score for a random subset (half) of the participants before computing the group-average difference value (cf. Aly et al., 2018). This procedure was repeated 1,000 times at each parcel to generate a null distribution of null difference values. As there were several parcels where the observed group differences in spatial ISC values exceeded all null values, *p*-values (one-tailed) were estimated for all parcels by using the null distribution’s mean and standard deviation to fit a normal distribution, allowing *p*-values to be calculated even when the observed differences in spatial ISC values for a parcel exceeded all null values. To correct for multiple comparisons across parcels, we controlled the False Discovery Rate (FDR) (Benjamini and Hochberg, 1995) using and FDR threshold of q= 0.05. The observed group-average difference value (i.e., without any flipped signs) was then compared against this null distribution. For the comparison Audio-Movies minus Backwards Speech, similar results are observed when stimulus duration is matched across conditions (Fig. S3).

#### Between-Group Comparisons

The group-average ISC value for the sighted participants was subtracted from the group-average spatial ISC value for the blind group, yielding a CB-S difference score. To determine statistical significance, a null distribution of 1,000 difference scores was generated by shuffling the labels of each subject’s group membership prior to computing each group-average ISC value, and then computing the CB-S difference score based on the randomly labeled data. This procedure was repeated 1,000 times at each parcel, and the true CB-S difference score (i.e., the resulting value when group membership labels were correctly assigned) was compared to this null distribution, yielding one *p*-value (one-tailed) per parcel. FDR correction for multiple comparisons (q= 0.05) was then applied to threshold the resulting *p*-values. To identify parcels where between-condition contrasts differ between groups, this procedure was performed on within-subject spatial ISC difference scores (e.g., Audio-Movies – Backwards Speech) rather than on the raw spatial ISC values themselves.

## Results

### Audio-Movies induce shared patterns in higher-cognitive regions in both groups, relative to Backwards Speech

Backwards Speech elicited reliable shared spatial patterns (spatial ISC) in auditory cortices across individuals for both blind and sighted groups. In addition, shared patterns were observed for the blind group in the right lateral and orbital frontal cortex, left precuneus and left superior extrastriate cortex. A direct comparison of the two groups did not reveal any parcels where spatial ISC values for the blind exceeded the sighted. There were no differences between the two groups in the visual cortices. The sighted group showed greater spatial ISC in left angular gyrus and left cingulate cortex, relative to the blind group (Fig. 2A). As expected, resting state stimuli did not elicit shared spatial patterns anywhere in the brain for either group.

**Figure 2.**
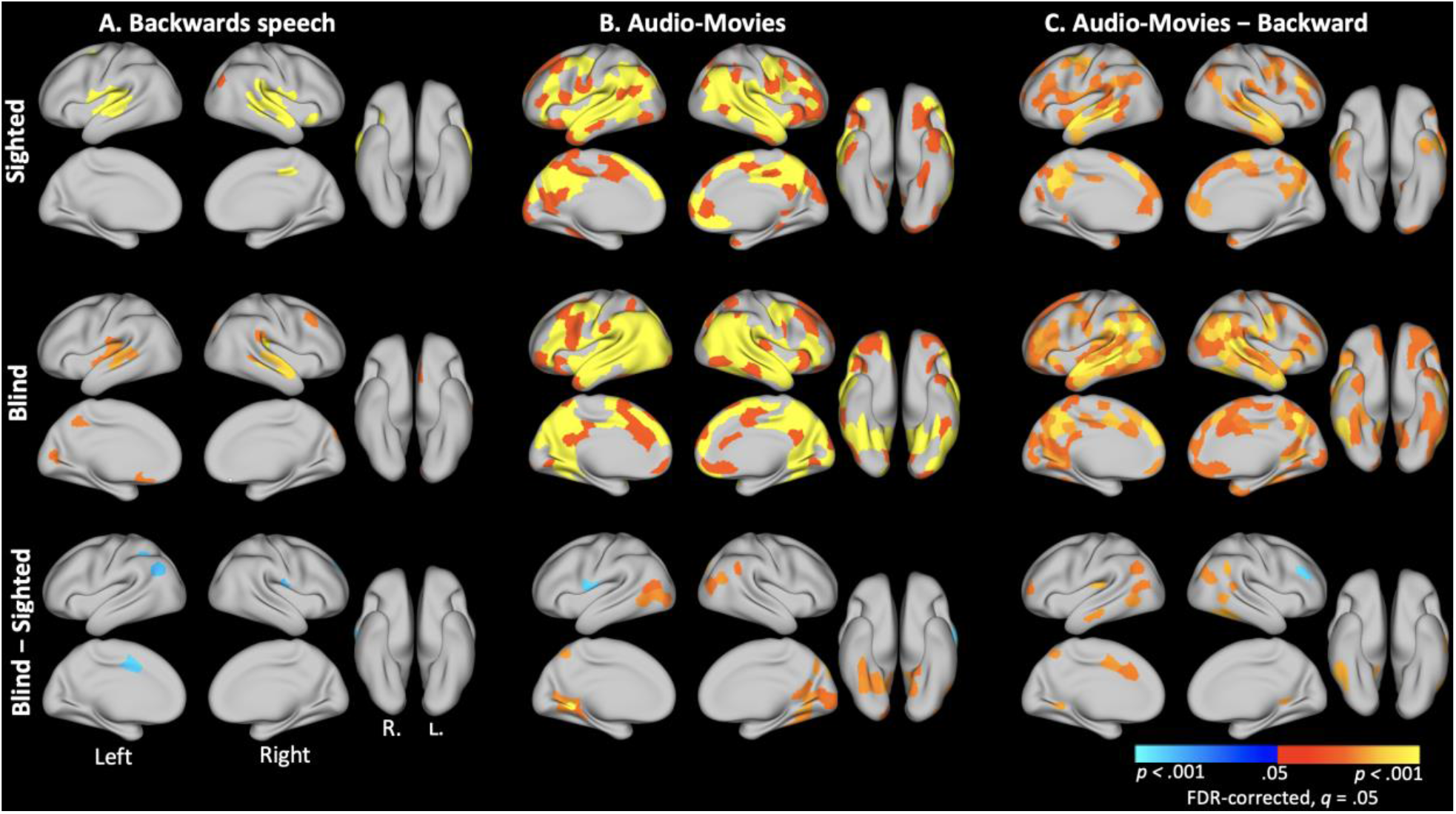
Parcels with reliable spatial ISC values both within and between subject groups for the (A) Backwards Speech condition, (B) the Movies condition, and (C) the contrast of Audio-Movies versus Backwards Speech. Maps for each individual stimulus condition are shown in Figures S2-S3. All maps corrected for multiple comparisons, *p*<0.05.

Audio-Movies elicited reliable shared spatial patterns not only in auditory cortices but also in higher-cognitive networks, including large segments of lateral temporal and prefrontal cortices, and the precuneus for both groups (Fig. 2B). For both blind and sighted groups, higher-cognitive networks showed reliably greater spatial ISC in response to the Audio-Movies as compared to Backwards Speech (Fig. 2C).

### Audio-Movies induce shared spatial patterns in visual cortices of blind participants

For the blind group, Audio-Movies elicited significant inter-subject spatial pattern correlations in occipital and occipito-temporal (OT) areas, including bilateral V1, bilateral lingual gyrus, right-lateralized posterior fusiform gyrus and inferior occipito-temporal cortex. In the sighted group, occipital involvement was less extensive. However, the part of occipital cortex directly ventral to the precuneus and one parcel in the left occipital pole, also showed reliable spatial ISC in the sighted group.

When blind and sighted groups were compared to each other directly, a subset of occipital regions with reliable spatial ISC for Audio-Movies in the blind group also showed significantly larger ISC in blind relative to sighted people. Spatial ISC was higher for Audio-Movies among blind people in ventral occipito-temporal cortex, right lingual gyrus, right OT cortex, and the right occipital pole, as well as medially in left V1 and V2 (Fig. 2B). Right OT cortex and bilateral medial lingual gyrus also showed a significant group-by-condition interaction: blind > sighted, Audio-Movies > Backwards Speech (Fig. 2C). In addition to these visual regions, spatial ISC increased for Audio-Movies versus Backwards Speech in the blind group, relative to the sighted, in the temporo-parietal junction bilaterally and left anterior cingulate cortex. The sighted group had larger spatial ISC than the blind group in one parcel in the right lateral prefrontal cortex. Spatial ISC maps for each individual stimulus condition are shown in Fig. S2.

A previously published analysis of these data tested for synchronized timecourses across subjects over the duration of each stimulus condition (i.e., temporal ISC instead of spatial ISC in the current study, cf. Loiotile et al., 2019). To compare the current spatial ISC results to the previous temporal ISC analysis with the same dataset, we plotted spatial ISC values (y-axis) as a function of temporal ISC values (x-axis) (Fig. 3) for Audio-Movies and Backwards Speech conditions and for each subject group. Parcels located in visual cortices are shown in red (Visual Network from Schaeffer et al. 2018 17-Network Parcellation). In the blind relative to the sighted group, the visual parcels shift right for the Audio-Movies condition, reflecting higher temporal ISC, and a subset of these visual parcels also shift upward, reflecting higher spatial ISC.

**Figure 3.**
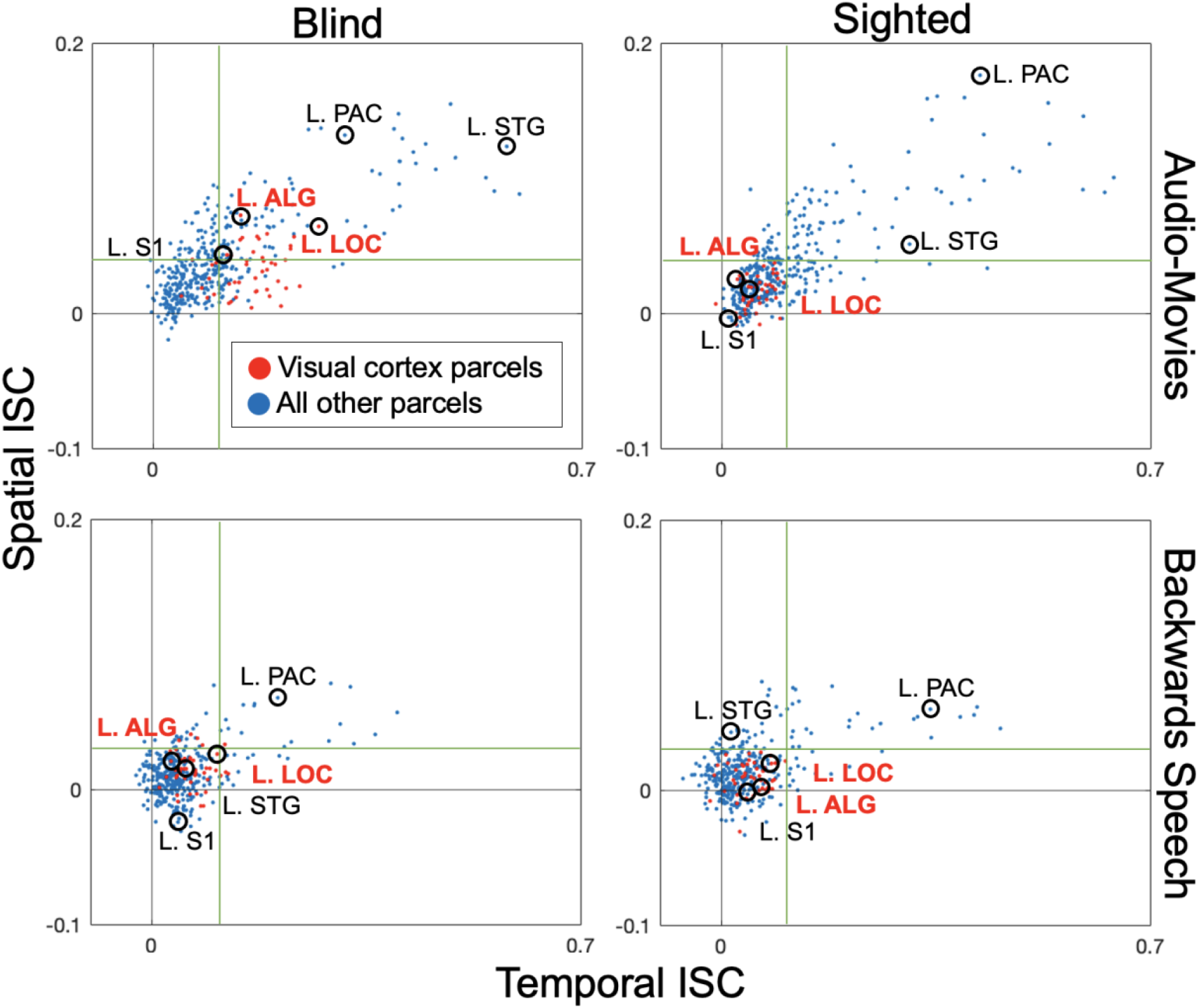
Scatterplots depicting parcel-level inter-subject correlation values (spatial ISC versus temporal ISC), separated by subject group and stimulus condition. Anatomical labels for example brain regions correspond to the nearest circled parcel dot. Red dots and region labels show parcels located in visual cortex. For each condition, the green lines delineate the maximum temporal ISC value (vertical line) and maximum spatial ISC value (horizontal line) observed in any visual cortex parcel in the sighted group. Spatial and temporal ISC values are more correlated in the Audio-Movies stimuli (Blind: *r*= 0.69; Sighted, *r*= 0.71) than in the Backwards Speech stimulus (Blind: *r*= 0.47; Sighted, *r*= 0.47). L= left; STG = superior temporal gyrus; PAC = primary auditory cortex; S1= primary somatosensory cortex; LOC = lateral occipital complex; ALG = anterior lingual gyrus

### Spoken narrative (Pie-Man) and sentences elicit shared spatial patterns in the visual cortices of blind but not sighted individuals

Results for the Pie-Man spoken narrative stimulus were qualitatively consistent with those observed in response to the Audio-Movie stimuli in both groups, however Pie-Man responses were weaker. In both groups, Pie-Man elicited shared spatial patterns in bilateral auditory cortex, bilateral medial prefrontal cortex, bilateral lateral temporal cortex, and precuneus. A direct comparison of the two groups revealed that Pie-Man elicited greater spatial ISC in the blind group in left lateral prefrontal cortex and parcels located in visual cortex, including bilateral medial visual cortex and right occipital pole (Fig. S4). Spatial ISC in right anterior temporal pole and left postcentral gyrus was greater among the sighted group.

The Sentence-Shuffle stimuli induced spatial ISC in similar areas in both groups, including bilateral auditory cortex and bilateral lateral temporal cortex. Blind people showed greater spatial ISC than the sighted for Sentence-Shuffle condition in left premotor cortex, right temporo-parietal junction, and one parcel in left medial visual cortex.

## Discussion

### Spatially consistent activity patterns emerge across visual cortices of different blind people

We observed segment-specific spatially distributed patterns of activity in the “visual” cortices of blind individuals in response to naturalistic auditory narratives. The spatial organization of these responses was similar across different blind individuals, such that direct comparisons of multi-voxel activity patterns across people revealed alignment in several regions of the occipital lobes, including medial and foveal V1 as well as regions in ventral occipitotemporal cortex. These findings suggest that information is spatially distributed within the visual cortices of different blind people in a consistent way. While we also observed some evidence of shared spatial patterns in the visual cortices of sighted individuals, the robustness and spatial extent of this effect was greater in the blind group.

### Spatial patterns of activity in “visual” cortex of blind participants reflect meaningful content

The shared spatial patterns in visual cortex selectively emerged in response to the meaningful auditory stimuli but not unintelligible backwards speech, indicating that these systematic activity patterns encode meaningful, high-level information in naturalistic narratives and not meaningless auditory input. The sound excerpts from movies contain several stimulus properties that the backwards speech stimuli lack. These naturalistic sound clips were expressly designed to entertain viewers with complex and inter-dependent events and characters. Unlike backwards speech, the movie sound excerpts feature plot lines that build over time, spoken language dialogues; music and emotive tones; environmental sounds and auditory cues to movement.

Although we do not know which aspects of the audio-movie contents are reflected in the spatial patterns of activity, the patterns are sensitive to information that varies over the duration of each movie excerpt. In order for a brain region to yield a statistically robust spatial ISC value, each audio-movie segment must, on average, show reliably greater pattern similarity across people for matching segments versus mismatching segments. Thus, not only are visual cortices sensitive to the difference between auditory movies and backwards speech, but they are also sensitive to differences among different segments of a given audio-movie.

Outside of visual cortex, several additional brain regions also showed consistent spatial patterns during meaningful movie excerpts relative to backwards speech in both the blind and sighted groups. Consistent with previous findings with sighted people, high spatial ISC emerged in several brain regions associated with higher-cognitive processing, including precuneus, medial frontal cortex, and bilaterally in the lateral frontal cortex and lateral temporal cortex. These areas are thought to support linguistic, semantic and social processes (Buckner et al., 2008; Hassabis et al., 2007; Binder et al., 2009; Fedorenko et al., 2011). Largely similar meaning-sensitive regions emerged across blind and sighted groups, consistent with previous evidence that lack of vision does not substantially alter the organization of the semantic system (e.g., Bedny et al., 2012; Hanjaras et al., 2017; Bottini et al., 2020). One possibility is that in blindness, the visual cortices assume functions similar to those performed in these higher-cognitive, non-visual regions in the sighted brain. However, more work is needed to determine the extent to which content is redundant across distinct brain regions that exhibit shared segment-level spatial patterns.

### Inter-subject synchrony exceeds spatial pattern alignment in the visual cortices of blind individuals

The current spatial ISC analysis enables the detection of shared “visual” cortex responses at a specific spatial scale (i.e., distributed across subsets of neighboring voxels) and at a particular grain of information (i.e., distinct ten-second segments within an audio-movie). Previous univariate analyses also show that “visual” cortices of different blind participants also aligned in temporal activity in response to audio-movie stimuli (i.e., temporal ISC) (Loiotile et al. 2019). The findings from these univariate temporal analysis are consistent with the multivariate spatial results we report here: shared temporal and spatial responses emerged in “visual” cortex, but more so in the blind group, and more for the meaningful auditory movies than for meaningless backwards speech.

Interestingly, although results of spatial and temporal ISC analysis generally align, spatial ISC effects observed in “visual” cortex of blind individuals are less anatomically widespread and robust than the temporal ISC effects (Fig. 3). Spatial ISC requires a greater degree of spatial (multivariate) alignment across people in order to detect effects, relative to temporal (univariate) ISC. One possible interpretation is that only a subset of the visual cortex regions that respond to auditory narratives contain maps of segment-specific information in individual people. Alternatively, despite general alignment, there may be greater variability in “visual” cortex information maps across different blind people, relative to visual cortices of sighted people and non-visual areas in both groups. Such variability could be related to lack of visual experience.

Some prior evidence supports the idea that cortical functional specialization is somewhat more variable across blind than sighted people. Previous studies find that laterality patterns vary to a greater extent across blind than sighted people (Lane et al., 2017; Abboud et al., 2019). This could contribute to lower alignment in visual cortex in the current study. Consistent with the idea of greater individual variability of cortical map across blind individuals, a recent MEG study revealed that semantic categories of auditory word stimuli were decodable from occipital cortex of blind individuals when analyses were performed at the individual-subjects level; however, cross-subject generalization was unreliable, indicating considerable variability in neural responses across blind individuals (Abboud et al., 2019). Together with prior findings, the present results provide both evidence for presence of reliable visual cortical maps across people who are blind and evidence that cortical alignment across people is somewhat reduced in blind relative to sighted people. One possible interpretation is that more consistent maps emerge when visual cortex assumes its evolutionarily predisposed function. A related possibility is that absence of vision leaves more room for variation during development, i.e., more free parameters. At present these hypotheses are highly speculative and require future testing.

### Future directions: Identify both shared and unique spatial patterns

In future work it will be important to apply similar analysis techniques to compare within and across participant alignment in blind and sighted populations by presenting the naturalistic stimuli with the same semantic content to the same blind individual across different testing sessions. This would allow quantification of alignment across sessions within blind and sighted people. Such designs would also enable better characterizing the spatial scale at which information is represented in “visual” cortices of individual blind adults. Previous work using multivariate analyses in individual participants revealed medial/lateral organization in the ventral stream and distinctive patterns within V1 (van den Hurk et al., 2017; Vetter et al., 2020). The current study used a between subject, naturalistic stimulus approach. Parcels within “visual” cortex that showed information sensitivity ranged in size from 1404 to 2790 mm^3^. In the current study, the spatial scale must be sufficiently coarse-grained to survive transformation from native space to a common brain template, yet fine-scale enough to differentiate between multiple distinct audio-narrative segments. Combining naturalistic stimulus and within-participant comparisons could reveal still more fine-scale representations that is sensitive to segment-specific patterns, reliable within a person but not between individuals (e.g., idiosyncratic organization).

In sum, the present results show that a common spatial organization for naturalistic auditory comprehension is present in the visual cortices of blind individuals. In this regard, deafferented “visual” cortices resemble high-level cortical areas typically associated with semantic, linguistic, and narrative processing. It is unlikely that “visual” cortices evolved representational maps specifically to represent auditory and semantic information. Nevertheless, existing anatomical constraints and shared non-visual experience among blind people is sufficient to produce significant alignment of cortical maps of visual cortex, paralleling the retinotopic organization observed in this region in the sighted brain.

## Acknowledgements

This work was supported by the National Institute of Health (NEI, R01 EY027352-01 to M.B.) and a National Science Foundation Postdoctoral Research Fellowship (SBE SPRF 1911650 to E.M.). We thank the blind and sighted individuals who participated in this study, and are grateful toward the blind community for its support of this research. We also thank the F. M. Kirby Research Center for Functional Brain Imaging at the Kennedy Krieger Institute.

## Supplementary Material

**Figure S1.**
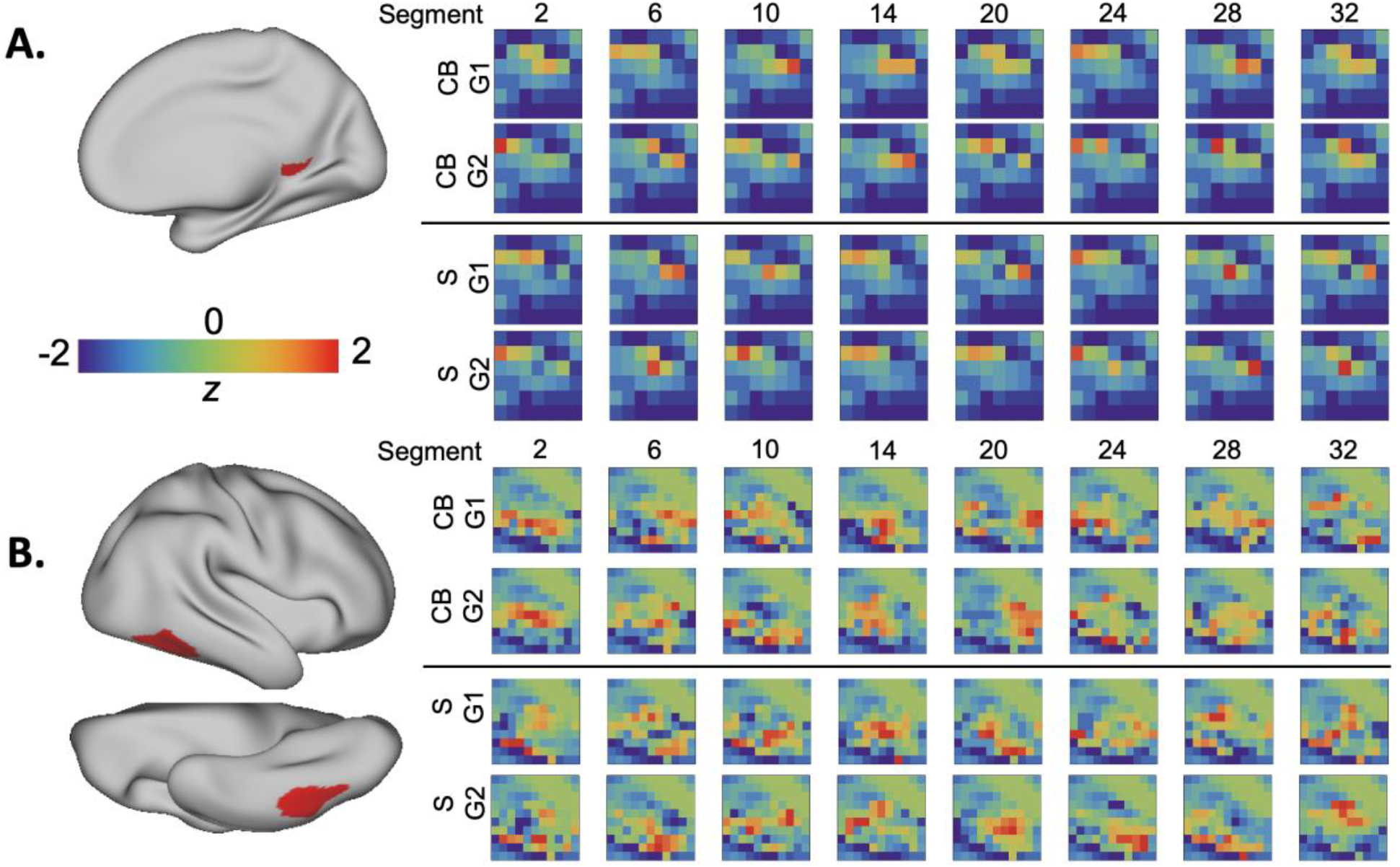
Segment-level pattern similarity between individuals. Squares depicting z-scored activity patterns response during individual segments show a single sagittal slice through each ROI. To visualize the spatial activity patterns for each segment, the data acquired during Movie 2 (Taken) were randomly split into four independent sub-groups and averaged across participants within each sub-group (CB group 1, n=11; CB group 2, n=11; S group 1, n=11, S group 2, n=10). The group mean TR-level patterns were then averaged across timepoints within a segment, yielding 32 segment patterns per sub-group. A. Segment patterns in a 52-voxel left hemisphere parcel labeled as “VisPeri_ExStrSup” (Visual peripheral, extrastriate superior cortex) in the Schaeffer et al. (2018) 17 Network Parcellation. Segment patterns depict a 42-voxel sagittal slice of the ROI. B. Segment patterns in a 223-voxel right hemisphere ROI, “DorsAttnA_TempOcc_2” (Dorsal attention, temporal occipital cortex). Segment patterns depict a 110-voxel slice of the ROI. CB = congenitally blind; S = sighted; G1 = group 1; G2 = group 2

**Figure S2.**
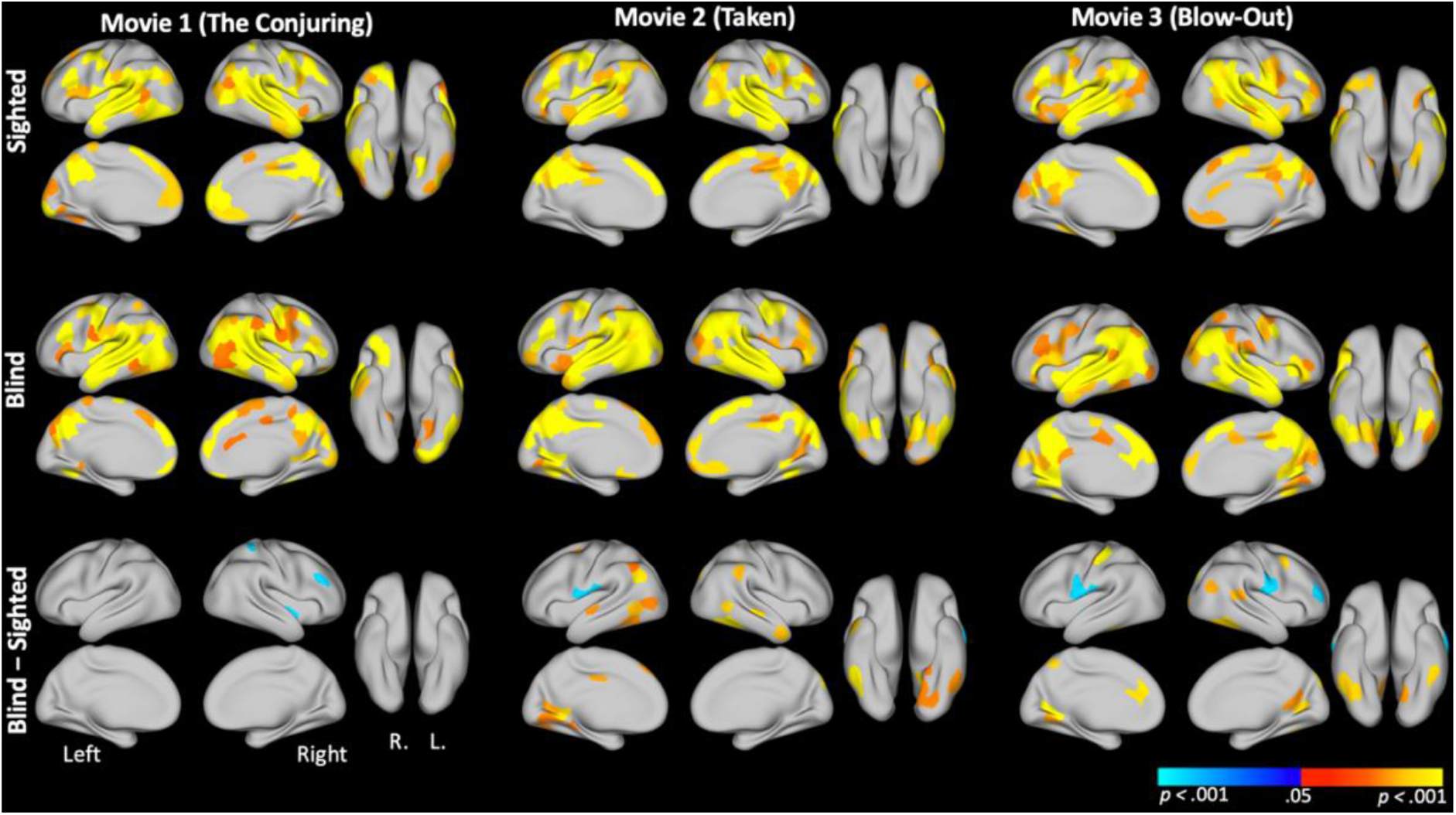
Spatial ISC parcel maps for each individual audio-movie stimulus. Within-group, within-condition comparisons (top two rows) were corrected for multiple comparisons using a bootstrap procedure to generate a null distribution of maximum correlation values (cf. Regev, Honey, Simony, & Hasson, 2013, *Journal of Neuroscience*). The between-group comparisons (third row) were corrected for multiple comparisons by applying FDR correction (q=0.05) to *p*-values which were computed by comparing observed group-level differences in spatial ISC values to null distribution in which group membership labels were shuffled one thousand times. All participant data were included in this analysis.

**Figure S3.**
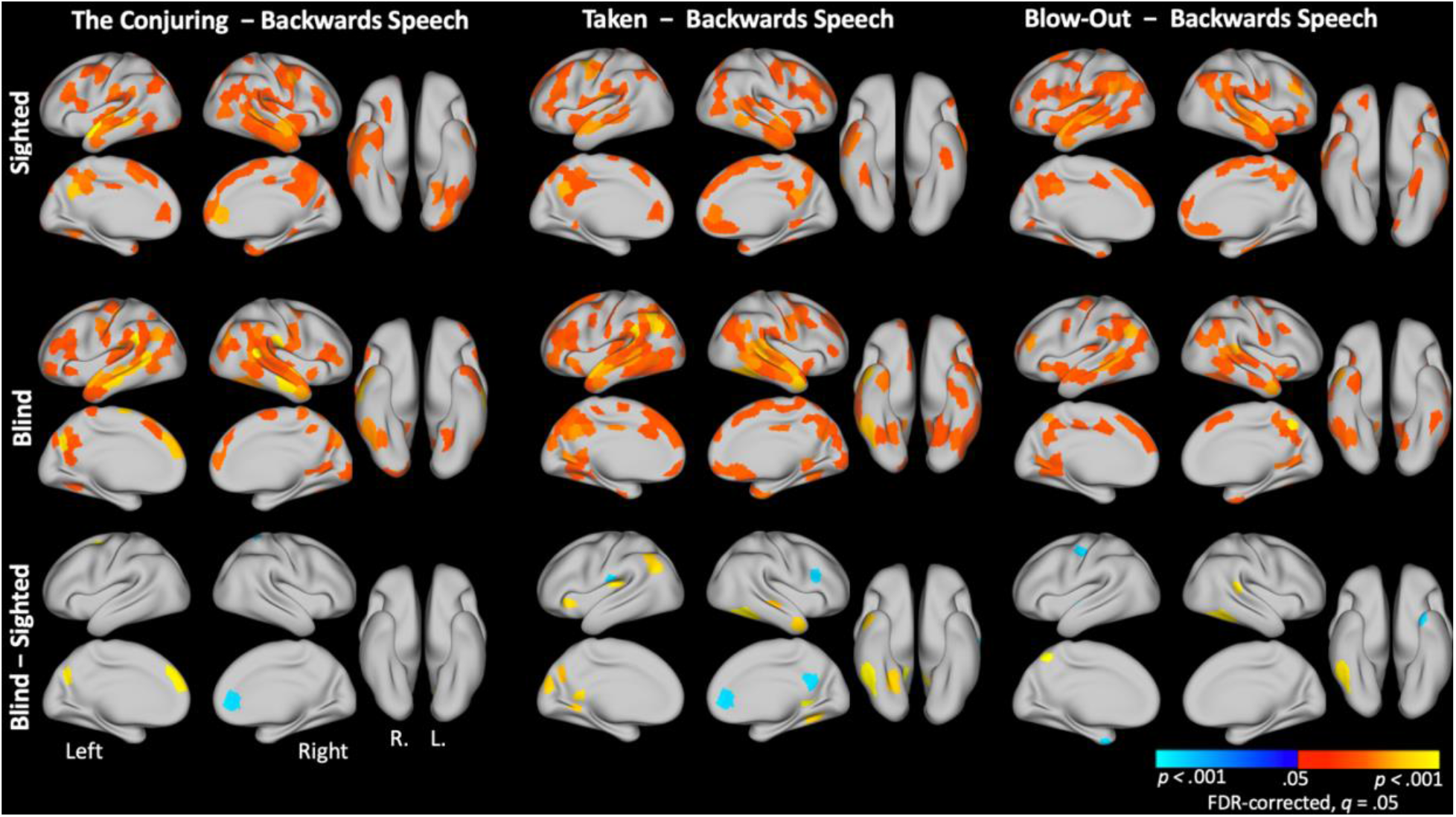
Spatial ISC parcel maps for each individual audio-movie stimulus minus backwards speech. Spatial ISC difference scores at every parcel were computed and then compared to a bootstrapped null distribution in which condition labels were shuffled one thousand times. Significance values were corrected for multiple comparisons with FDR correction (q=0.05). All participant data were included in this analysis.

**Figure S4.**
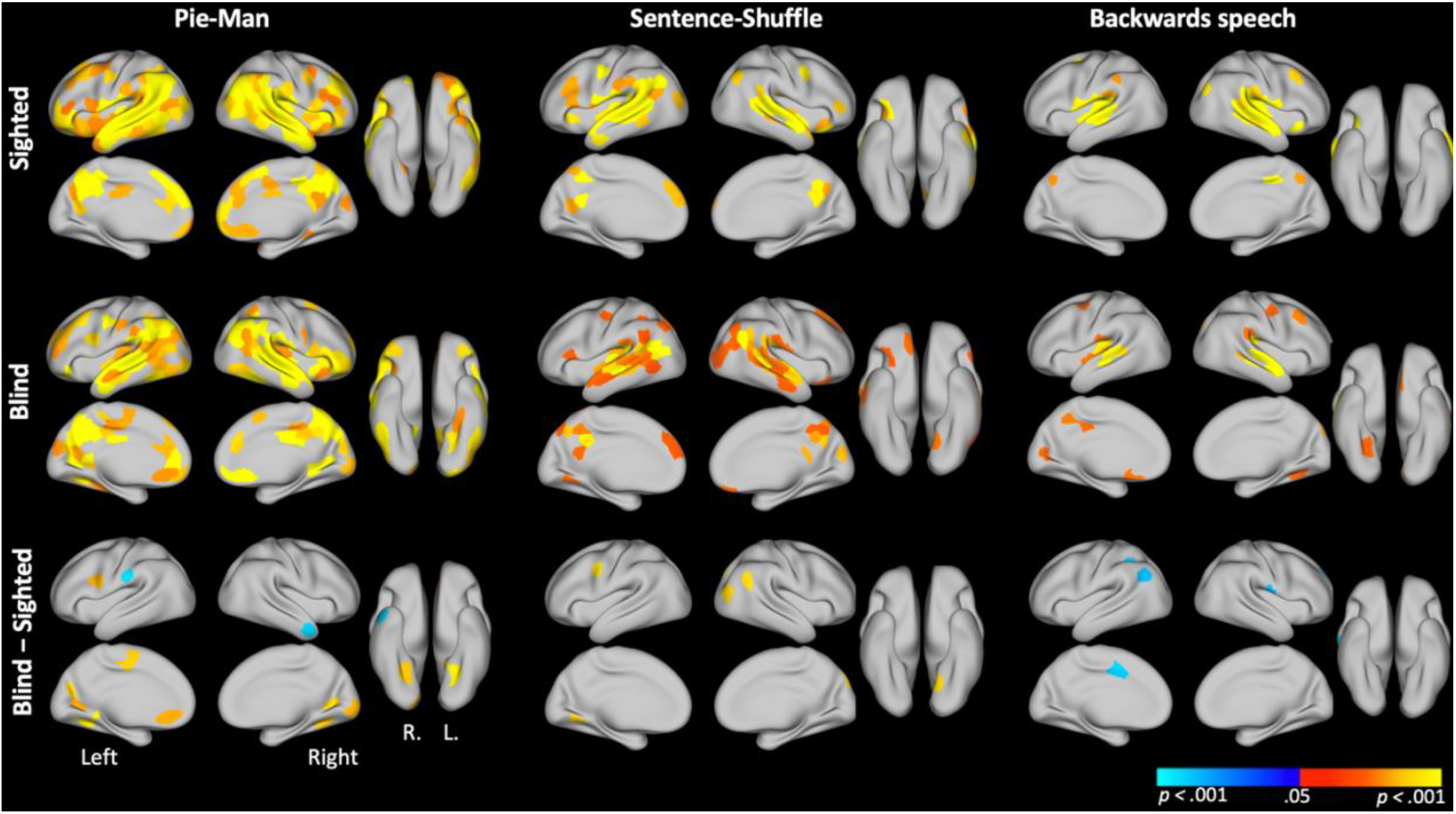
Spatial ISC parcel maps for intact and degraded versions of the Pie-Man stimuli. Within-group, within-condition thresholding (first two rows) was applied using the procedures described for Figure S1. Between-group thresholds (third row) were determined via bootstrapping by shuffling group membership labels one thousand times to generate a null distribution, then FDR correction was applied (q=0.05). All available participant data were included in this analysis.

**Figure S5.**
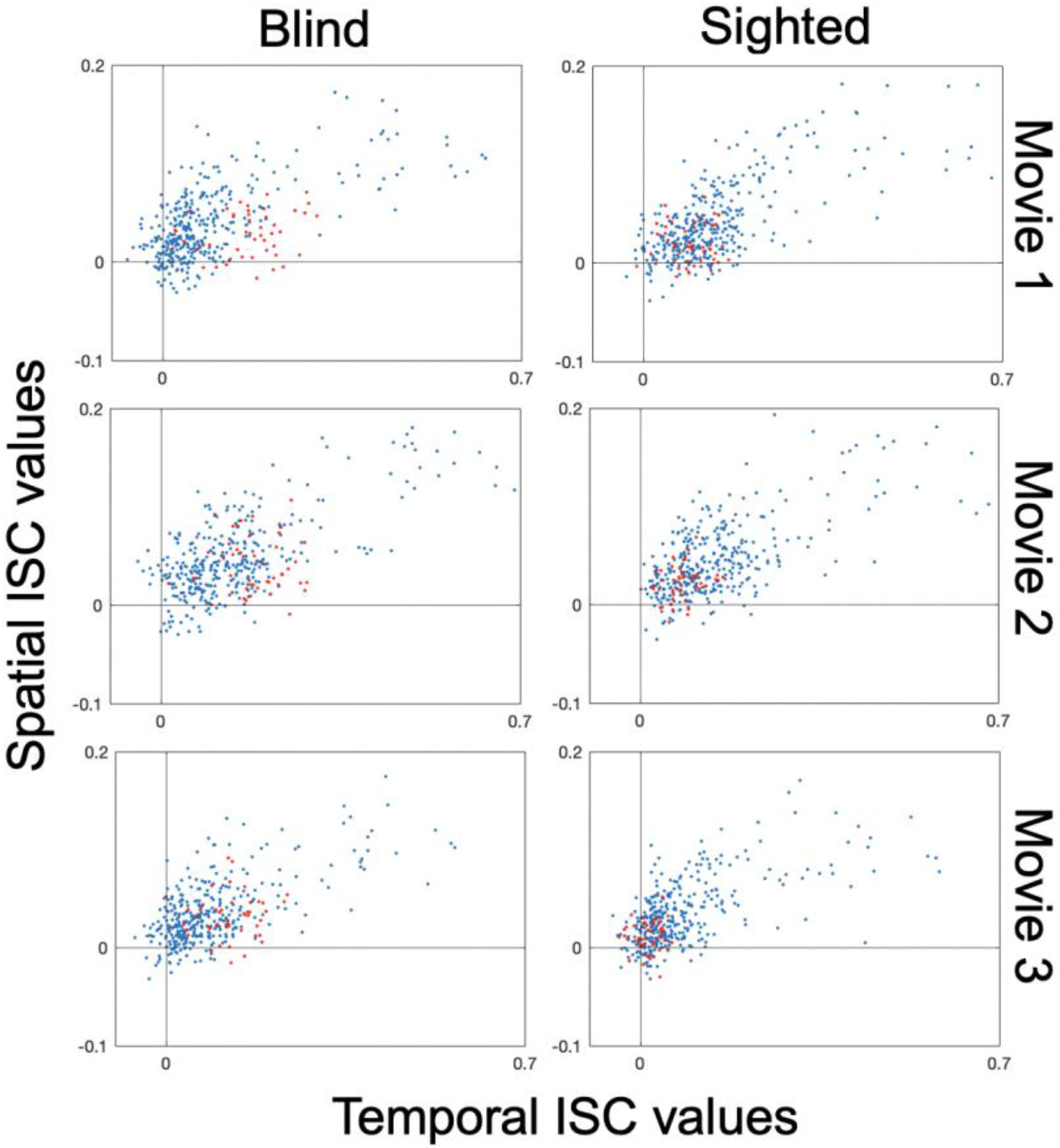
Relationship between spatial ISC and temporal ISC in each brain parcel, for each individual movie stimuli. Red dots depict values for parcels located in the Visual Network as identified in the Schaeffer et al. (2018) 17-Network brain parcellation.

**Figure S6.**
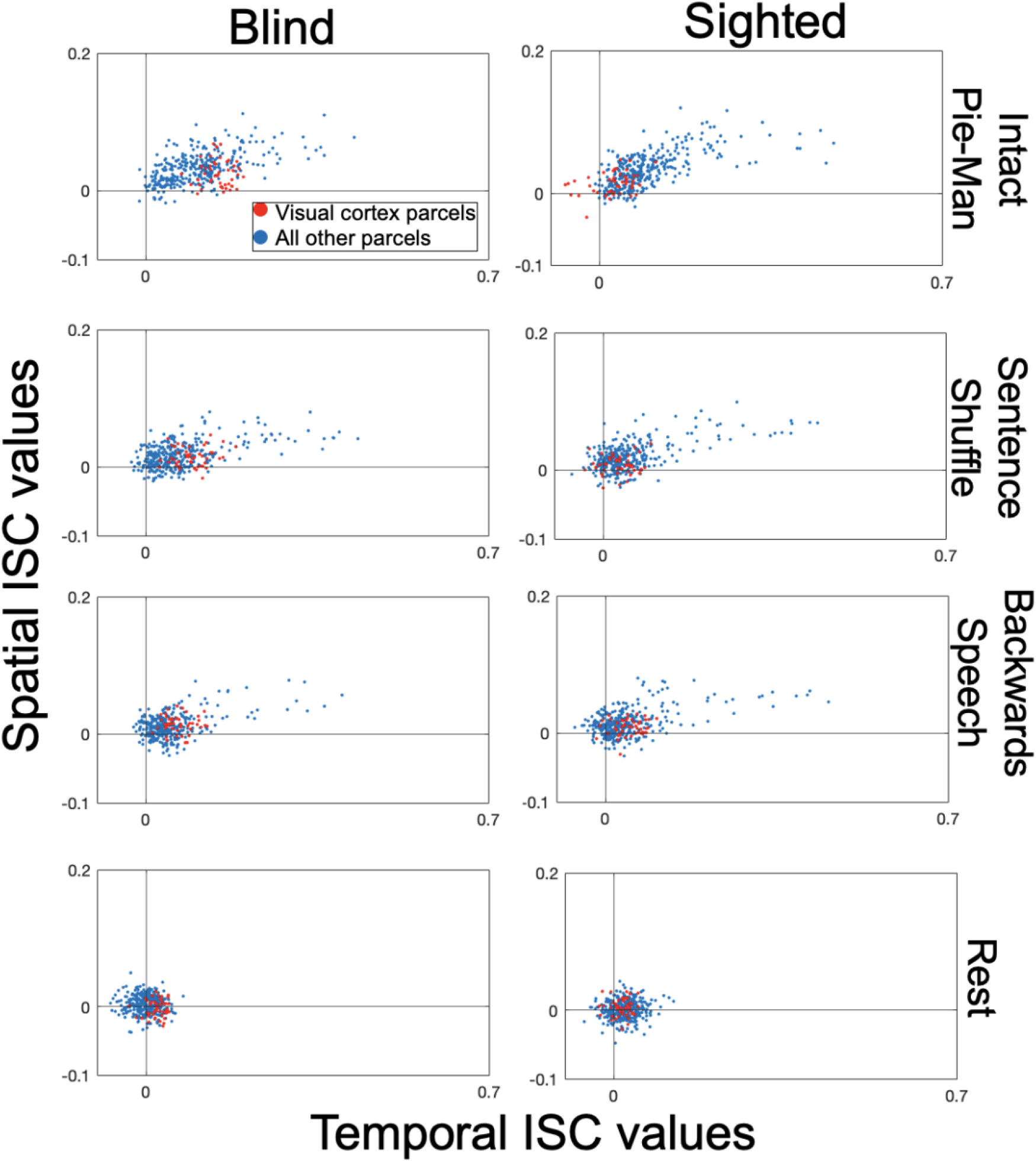
Relationship between spatial ISC and temporal ISC in each brain parcel, for the Rest condition and Pie-Man stimuli. Red dots depict values for Visual Network parcels.

**Figure S7.**
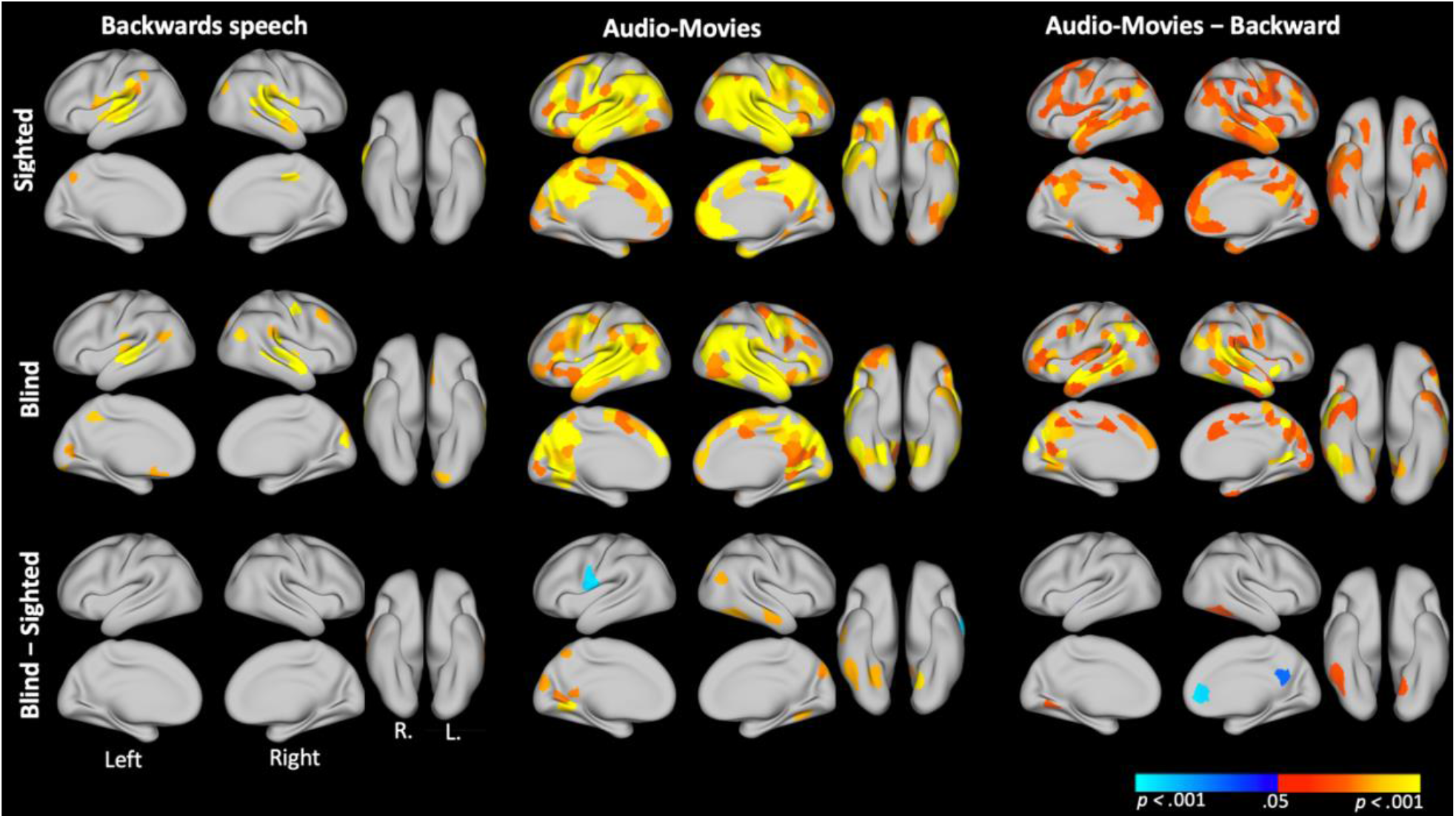
Spatial ISC parcel maps for the concatenated audio-movie stimuli versus backwards speech, limited to participants with high accuracy scores (3/5 or better) on each of the post-scan multiple-choice comprehension tests for the three movies (CB n= 13; S n= 15). Thresholding procedures follow those described for Figure S1.

**Figure S8.**
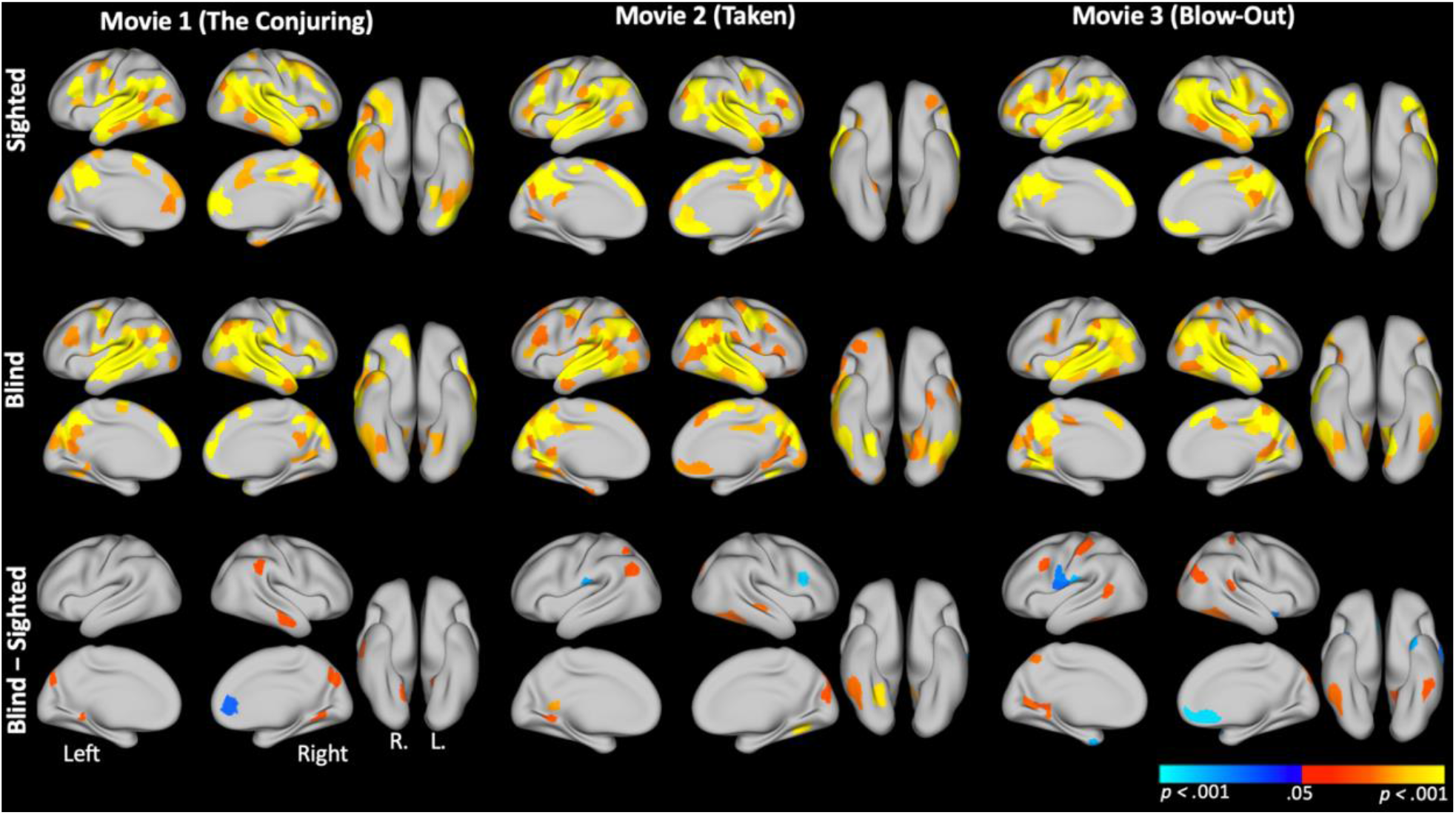
Spatial ISC parcel maps for each individual audio-movie, limited to participants with high accuracy scores (3/5 or better) on the corresponding post-scan multiple choice comprehension test for each respective condition (Movie 1: CB n= 15, S n= 18; Movie 2: CB n= 17, S n= 16; Movie 3: CB n= 17, S n= 17). Thresholding procedures follow those described for Figure S1.

